# The Host Cell ViroCheckpoint: Identification and Pharmacologic Targeting of Novel Mechanistic Determinants of Coronavirus-Mediated Hijacked Cell States

**DOI:** 10.1101/2020.05.12.091256

**Authors:** Pasquale Laise, Gideon Bosker, Xiaoyun Sun, Yao Shen, Eugene F. Douglass, Charles Karan, Ronald B. Realubit, Sergey Pampou, Andrea Califano, Mariano J. Alvarez

**Affiliations:** DarwinHealth Inc, New York, NY, USA; Department of Systems Biology, Columbia University Irving Medical Center, New York, NY, USA; Herbert Irving Comprehensive Cancer Center, Columbia University Irving Medical Center, New York, NY, USA; Department of Medicine, Columbia University Irving Medical Center, New York, NY, USA; Department of Biochemistry & Molecular Biophysics, Columbia University Irving Medical Center, New York, NY, USA; Department of Biomedical Informatics, Columbia University Irving Medical Center, New York, NY, USA

**Keywords:** Coronavirus, Regulatory networks, Master regulator, Anti-viral drugs

## Abstract

Most antiviral agents are designed to target virus-specific proteins and mechanisms rather than the host cell proteins that are critically dysregulated following virus-mediated reprogramming of the host cell transcriptional state. To overcome these limitations, we propose that elucidation and pharmacologic targeting of host cell Master Regulator proteins—whose aberrant activities govern the reprogramed state of infected-coronavirus cells—presents unique opportunities to develop novel mechanism-based therapeutic approaches to antiviral therapy, either as monotherapy or as a complement to established treatments. Specifically, we propose that a small module of host cell Master Regulator proteins (ViroCheckpoint) is hijacked by the virus to support its efficient replication and release. Conventional methodologies are not well suited to elucidate these potentially targetable proteins. By using the VIPER network-based algorithm, we successfully interrogated 12h, 24h, and 48h signatures from Calu-3 lung adenocarcinoma cells infected with SARS-CoV, to elucidate the time-dependent reprogramming of host cells and associated Master Regulator proteins. We used the NYS CLIA-certified Darwin OncoTreat algorithm, with an existing database of RNASeq profiles following cell perturbation with 133 FDA-approved and 195 late-stage experimental compounds, to identify drugs capable of virtually abrogating the virus-induced Master Regulator signature. This approach to drug prioritization and repurposing can be trivially extended to other viral pathogens, including SARS-CoV-2, as soon as the relevant infection signature becomes available.

## Introduction

SARS-CoV is an enveloped, positive-sense, single-stranded RNA virus of the genera *Betacoronavirus* introduced into the human population from an animal reservoir and culminating in a lethal epidemic in 2002-03, affecting 8,098 individuals, 774 of whom died (9.6%)(1). The virus shares 79% genome sequence identity with SARS-CoV-2, which is responsible for the current COVID-19 pandemic(2). SARS-CoV can generate a rapid inflammatory cascade eventually leading to pneumonia or severe acute respiratory syndrome (SARS), characterized by diffuse alveolar damage, extensive disruption of epithelial cells and accumulation of reactive macrophages(3). Similar to SARS-CoV-2, SARS-CoV spike protein S binds to angiotensin converting enzyme 2 (ACE2), which is widely expressed on the cell membrane of oral, lung, and nasal mucosa epithelial cells, arterial smooth muscle and venous endothelial cells, as well of other organs, including stomach, small intestine, colon, skin, lymph nodes, spleen, liver, kidney, and brain(4). Supportive care—including prevention of Acute Respiratory Distress Syndrome (ARDS), multi-organ failure, and secondary infections—remains the foundational approach for managing serious infections caused by coronaviruses, although preliminary analysis of a recently-reported, prospective, randomized, placebo-controlled trial, suggests that patients receiving remdesivir recovered faster than those receiving placebo(5–7). Despite early optimism and approval on May 1^st^, 2020 of remdesivir for emergency use in hospitalized patients with COVID-19, no other specific antiviral treatment has been proven to be effective in randomized, placebo-controlled trials(5, 6). Consequently, there remains a formidable unmet need to identify pharmacologic treatments, alone or in combination—directly targeting either viral mechanisms and/or host cell factors—that significantly inhibit viral replication and, by extension, minimize progression of target organ failure associated with COVID-19.

Current efforts focusing on antiviral drug discovery can be summarized as belonging to two broad strategies: (a) disrupting the synthesis and assembly of viral proteins or (b) targeting host proteins and mechanisms required by the viral replication cycle. The first strategy has yielded drugs targeting (i) viral proteases, required for processing of the virus large replicase polyprotein 1a, producing non-structural proteins involved in viral transcription and replication(5, 8); (ii) RNA-dependent RNA-polymerase, using guanosine and adenosine analogs, as well as acyclovir derivatives; (iii) virus helicases; (iv) viral spike proteins, with antibodies, peptide decoys and carbohydrate-binding agents; and (v) structural proteins such as those maintaining ion channel activity of CoV E protein and RNA-binding affinity of CoV N protein(5, 6, 9, 10). Although virus-targeting approaches have the advantage of being specific, and, therefore, generally offer acceptable toxicity profiles, targeting viral products typically restricts the applicability of antiviral agents to only one, or only a few, closely related virus species. Moreover, due to the high mutation rate of viral genomes, such drugs are prone to rapid virus adaptation by resistant strain selection(11, 12). Considering the time required to develop new pharmacologic agents, this strategy has proven unsuitable to address new viral epidemics and pandemics in real time.

In contrast, targeting host cell proteins, especially at an early stage when viral hijacking of host mechanisms may still be reversible, may have more universal and longer term value because the same host factors may be required by multiple, potentially unrelated viral species and because host target proteins mutate far less rapidly than viral proteins, thereby limiting emergence of drug resistance(13). Unfortunately, pharmacologic targeting of host factors is more commonly associated with toxicity, thereby limiting clinical application of many drugs identified as potential anti-viral agents *in vitro*, for instance, with anti-CoV drugs *EC*_50_ markedly exceeding their maximum tolerated serum concentration (*C_max_*)(5). Despite these translational challenges, current approaches to target host proteins are primarily based on either boosting innate anti-viral immune response, in particular interferon response, or targeting proteins and processes mediating viral infection, such as ACE2 receptors(14), cell surface and endosomal proteases(15), and clathrin mediated endocytosis(16). Moreover, broad availability of high-throughput screening approaches has allowed the purposing and repurposing of drugs based on their effect on virus replication(16–19), leading to identification of several anti-coronavirus candidates, such as chloroquine, tamoxifen, dasatinib and lopinavir, among others(16, 19). Yet, this approach is limited by the idiosyncratic nature of the *in vitro* models used in antiviral screens and by drug concentrations that may not be achievable in patients(5).

More recently, systems biology approaches, including temporal kinome analysis(20) and proteomics(21–24), have also been used to identify protein kinases—and associated pathways—modulated in response to virus infection, as well as to generate virus-host protein-protein interactomes (PPI). These methods also present an opportunity to develop and test host-targeting therapeutic approaches that apply functional genomics to the “infected system as a whole.”(24) The output of these predictions can be used to direct drug repurposing efforts(21–23) and to design more focused *in vitro* screens, with models that better recapitulate disease pathophysiology, such as primary cells, organoids or 3D organ-on-chip systems(25).

Coronaviruses have been shown to extensively hijack the cellular machinery of host cells they infect; as one example, this class of viruses induces arrest in S phase, allowing them to benefit from physiological alterations they induce in host cells that enhance their reproductive rate(26). As shown for other physiologic(27–29) and pathologic cell states—among them, cancer(30–34), neurodegeneration(35, 36), and diabetes(29)—we propose that such transcriptionally “locked” states are established by the virus and maintained by a handful of Master Regulator (MR) proteins, organized within a highly auto-regulated protein module, or checkpoint (see Califano & Alvarez(30) for a recent perspective). For simplicity, in a viral infection context, we will call such modules “ViroCheckpoints.” Accordingly, we propose that aberrant, virus-mediated activation of a ViroCheckpoint is ultimately responsible for creating a transcriptionally “locked” cellular context that is primed for viral replication and release. We thus propose ViroCheckpoint activity reversal as a potentially valuable therapeutic strategy for pharmacologic intervention.

Here we show that time-dependent, SARS-CoV-mediated ViroCheckpoints—and the specific MR proteins of which they are comprised—can be effectively elucidated by network-based analysis using the Virtual Inference of Protein activity by Enriched Regulon (VIPER) algorithm(37). More importantly, once the MR protein identity is available, drugs can be effectively and reproducibly prioritized based on their ability to invert the activity of ViroCheckpoint MR proteins, using the OncoTreat algorithm(34), a NYS CLIA-certified algorithm that is used routinely on cancer patients at Columbia University.(38)

Accurate identification of virus-dependent MR proteins permits deployment of the same OncoTreat-based methodological approach for mechanism-based repurposing or development of new drugs with potential anti-viral activity. To avoid confusion, we will use the term “ViroTreat” to indicate the virus-specific version of OncoTreat. Specifically, ViroTreat uses the full repertoire of virus-induced MR proteins in the ViroCheckpoint as a reporter assay to identify drugs capable of reversing its activity(34), thereby preventing emergence of or abrogating the virus-mediated transcriptional locked state. While limited by the availability of data on SARS-CoV-2, including of infection in an appropriate pathophysiologic cell context, we provide proof of concept that this approach can be applied to prioritizing FDA-approved and late-stage investigational drugs representing potential antiviral agents for SARS-CoV based on infection in cancer-related lung epithelial cells.

## Results

### Elucidating MRs of SARS-CoV infection in lung epithelial cells

To identify candidate MR proteins that mechanistically regulate the host cell gene expression signature induced by SARS-CoV infection (i.e. the SARS-CoV ViroCheckpoint), we applied the VIPER algorithm to a previously-published, microarray-based gene expression signature of a Calu-3 lung adenocarcinoma cell clone expressing elevated ACE2 levels, compared to the parental line, at 12h, 24h, and 48h following infection with SARS-CoV at MOI = 0.1(39). A total of 6,054 regulatory proteins were considered in the analysis, including 1,793 transcription factors (TFs), 656 co-transcription factors (co-TFs), and 3,755 signaling proteins (SP).

Similar to a highly-multiplexed gene reporter assay, VIPER measures the activity of an individual protein based on the enrichment of its positively regulated and repressed targets in genes that are over- and under-expressed in a specific cell state, compared to a control(37). We have shown that VIPER can accurately measure the activity of >70% of regulatory proteins and, as a result, the algorithm has been used to elucidate MRs of both pathologic(31–33, 35, 36, 40, 41) and physiologic cell states(27–29) that have been experimentally validated. Moreover, VIPER-inferred protein activity has been shown to provide a better biomarker of cell phenotype than the original transcriptional profile(30, 34, 42, 43); and, importantly, is a better reporter for validating clinically relevant drug sensitivity(44). Accordingly, VIPER requires a differential expression signature for each sample to be analyzed and a regulatory model comprising the transcriptional targets of each regulatory protein. For the former, we computed a differential gene expression signature for each SARS-CoV infected sample, by comparing it to three 12h mock control replicates. For the latter, we leveraged a transcriptional regulatory model (interactome) generated by ARACNe(45) analysis of 517 samples in the lung adenocarcinoma cohort of The Cancer Genome Atlas (TCGA)(37). Use of a cancer-related interactome is well justified as we have shown that protein transcriptional targets are highly conserved between cancer and normal cells(28).

The analysis revealed *n* = 236 proteins, whose activity was significantly affected by SARS-CoV infection in at least one time point (*p* < 10^−5^, Bonferroni Corrected (BC), see Supplementary Table 1). Examination of the top 10 activated MR proteins at each of the evaluated time-points (Fig. 1a) revealed the presence of canonical cell-cycle regulators, including (a) cyclins (CCNA2), and other proteins involved in G1/S transition(46) (E2F8 and UHRF1); (b) S-phase proteins, such as topoisomerases (TOP2A(47)) and other factors involved in S-phase cell cycle arrest(48) (CHEK1, GTSE1); (c) mitotic checkpoint proteins(49) (BUB1B, KIF11 and NDC80); and (d) proteins involved in nucleotide synthesis (GMPS). These showed significant activation as early as 12h after SARS-CoV infection. In contrast, established innate immune response proteins were also found among the top activated MRs, including IFN-induced factors(50) (MX1, IRF9 and IFI27) but their activation became most evident only at the latest time point (48h). Interestingly, some proteins previously identified as key tumor MRs were strongly activated, such as FOXM1 and CENPF(33, 51), although this may be a byproduct of the cancer related nature of the Calu-3 cells used in the infection assays.

**Fig. 1.**
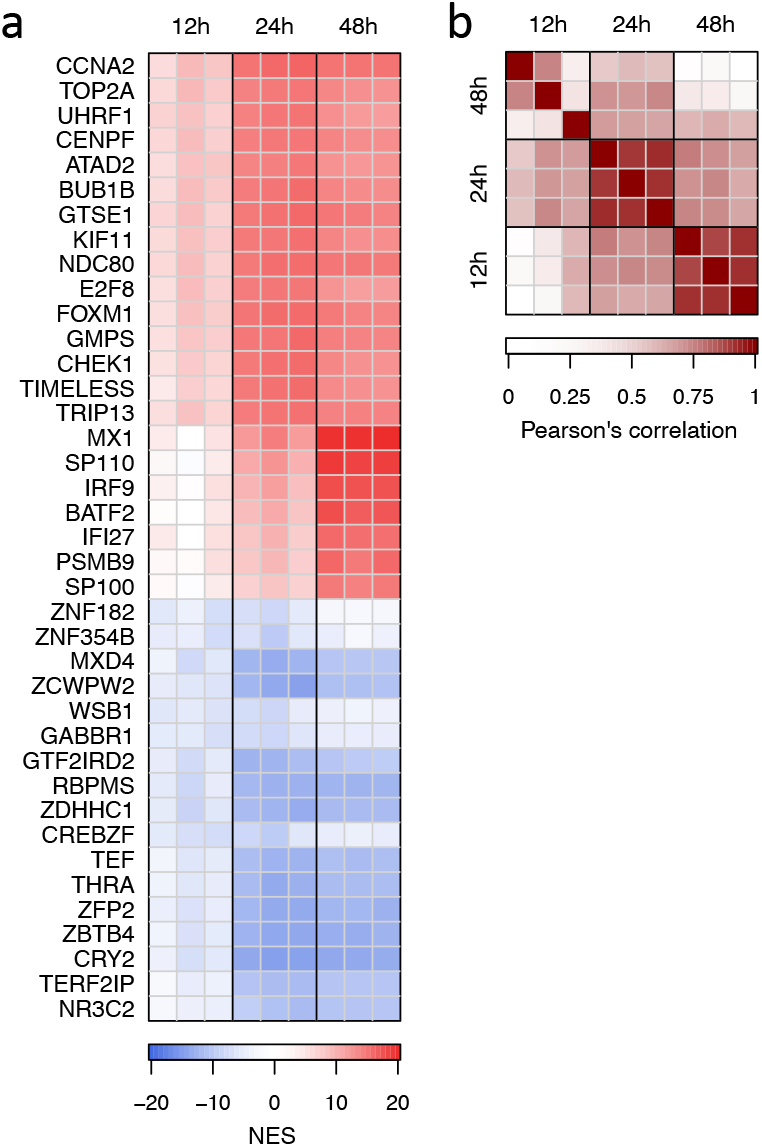
SARS-CoV-induced ViroCheckpoint in Calu-3 lung adenocarcinoma cells. Heatmap showing the VIPER-inferred protein activity, expressed as normalized enrichment score (NES), for the top 10 most activated and the top 10 most inactivated proteins in response to SARS-CoV infection for each of the three time points. Heatmap showing the similarity between the SARS-CoV induced protein activity signatures, expressed as Pearson’s correlation coefficient.

We then systematically evaluated whether viral infection could affect host proteins known to be involved in SARS-CoV host-pathogen protein-protein interactions (PPI). We based this analysis on a set of 36 proteins previously identified by high-throughput yeast-2-hybrid screen and validated by luciferase assays(23). Of the 36, 12 were represented among our set of 6,054 regulons and could thus be assessed for enrichment in SARS-CoV-induced differentially active proteins. Despite the low statistical power of a test based on only 12 proteins, enrichment was statistically significant for the 12h activity signature (*p* < 0.01, Supplementary Fig. 1a). Enrichment was borderline non-significant at 24h (*p* = 0.08), and not significant at 48h (Supplementary Figs. 1b and c).

To increase the test’s sensitivity, we leveraged a larger set of proteins identified as PPI for 26 of the 29 proteins coded by the closely related SARS-CoV-2 virus, as identified by massspec analysis of pull-down assays(21). Of 332 host proteins identified by that analysis, 89 were represented among those analyzed by VIPER. Confirming the prior results, enrichment was highly significant (*p*_12*h*_ < 10^−5^ by 2-tail aREA test(37); *p*_24*h*_ < 0.01 and *p*_48*h*_ < 0.001 by 1-tail aREA test, see Supplementary Fig. 1g, k and l, respectively). Interestingly, while enrichment was significant at all three time points, (*p* < 0.01, 1-tail aREA test, Supplementary Fig. 1j–l), several of the human SARS-CoV-2 PPIs activated at 12h became inactivated at later time points (Supplementary Fig. 1h–i).

Correlation analysis showed a gradual shift in protein-activity signatures from 12h to 48h after infection (Fig. 1b), suggesting dynamic activation and inactivation of a diverse repertoire of genetic programs by virus-host interaction and thus dynamic transition across multiple, time-dependent ViroCheckpoints. To gain insight into the biological programs most profoundly affected by SARS-CoV infection, we performed Gene-Set Enrichment Analysis (GSEA)(52) of a set of 50 biologically-relevant hallmark gene-sets from MSigDB(53) in differentially active, infection-mediated proteins (Fig. 2). The analysis identified four time-dependent program classes including: (a) cell cycle programs, consistently up-regulated at all three time points; (b) immune-related programs, associated with interferon response, inflammatory response, TNF-*α*, and IL-6/JACK/STAT3 signaling, which were progressively upregulated over time; (c) DNA repair pathways and (d) PI3K/AKT/mTOR programs more strongly activated at 12h (Fig. 2).

**Fig. 2.**
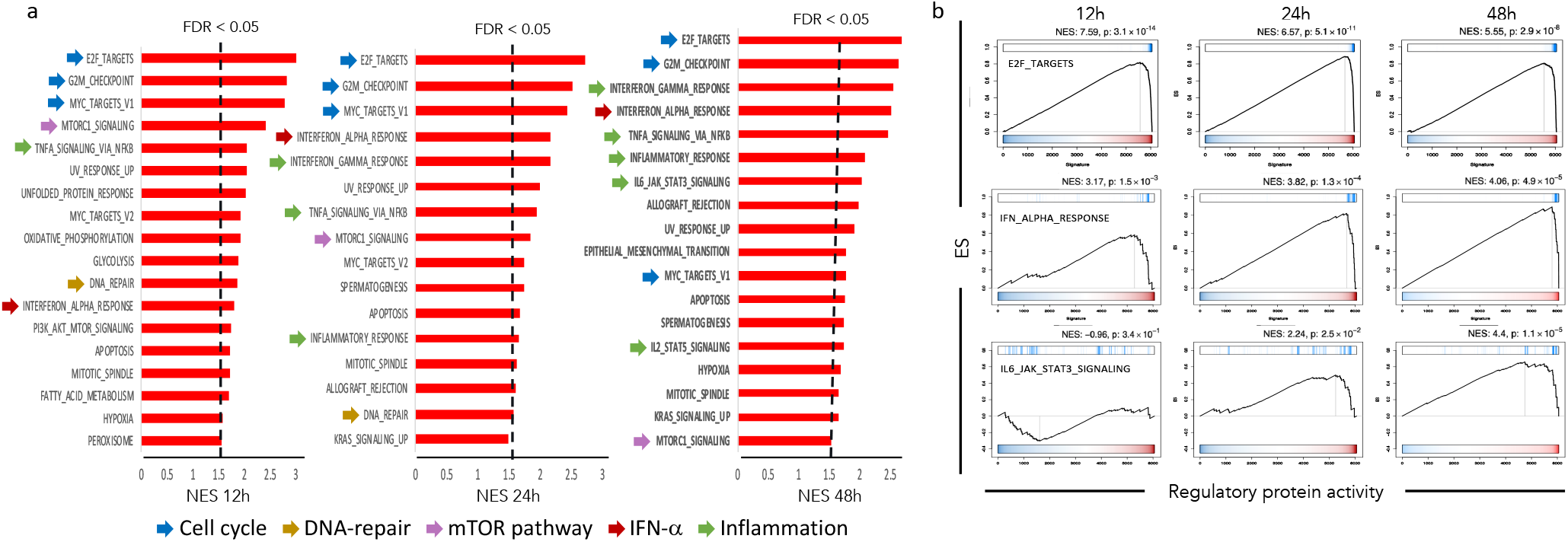
Biological programs activated by SARS-CoV infection. (a) Hallmark gene-sets from MSigDB significantly enriched (FDR < 0.05) in proteins activated at 12h, 24h and 48h after SARS-CoV infection. The bars indicate the GSEA-estimated Normalized Enrichment Score (NES). Pathways and processes related to cell cycle progression and cell proliferation, DNA-repair, mTOR, IFN-*α* and inflammation are indicated by blue, yellow, purple, red and green arrows, respectively. (b) GSEA plots showing the enrichment of E2F-targets, IFN-*α*-response and IL6/JAK/STAT pathway hallmark gene-sets on the differential activity of 6,054 regulatory proteins at 12h, 24h and 48h after SARS-CoV infection. The x-axis shows the regulatory proteins sorted from the most inactivated (left), to the most activated (right) in response to viral infection. The y-axis shows the enrichment score estimated by GSEA. The blue vertical lines indicate the proteins annotated as part of each of the analyzed biological programs/pathways.

Consistent with the multifarious effects that coronaviruses are known to exert through their complex, synchronized modulations of cell cycle progression, interferon antagonism, interleukin 6 and 8 induction, and host protein synthesis(26), these findings disclose a time-dependency, with early vs. late activation of protein signatures each linked to a distinct set of biofunctional hallmarks resulting from a virus-governed reconfiguration of the host cell’s regulatory state, with alterations in cell cycle during the initial post-infection phase, followed by a phase characterized by ignition of pro-inflammatory cytokine signaling pathways.

### ViroTreat analysis of SARS-CoV infected cells identifies novel therapeutic targets for drug repurposing

We have previously developed and validated a systematic approach (OncoTreat) for identifying drugs and compounds capable of reversing the aberrant activity of all Tumor Checkpoint MRs, representing mechanistic determinants of cell state, on a patient by patient basis(34). As a direct result of the high reproducibility demonstrated by VIPER,(37) the test has been certified by the NYS-CLIA laboratory and is available in the United States from the Columbia University Laboratory of Personalized Genomic Medicine(38); and, in China, from the Xiamen Encheng Group Ltd.

OncoTreat is used routinely to assess potential therapy for cancer patients who are progressing on standard of care, as part of the Columbia Precision Oncology Initiative(54). Despite the fact that it was originally developed for deployment and drug prioritization in the setting of precision oncology, the OncoTreat methodology is fully generalizable and can be applied to any state transition and any drug collection, including transitions related to and induced by viral infection. To avoid confusion, we will use the term ViroTreat to refer to the algorithm when used to identify antiviral drugs (see description in Fig. 3).

**Fig. 3.**
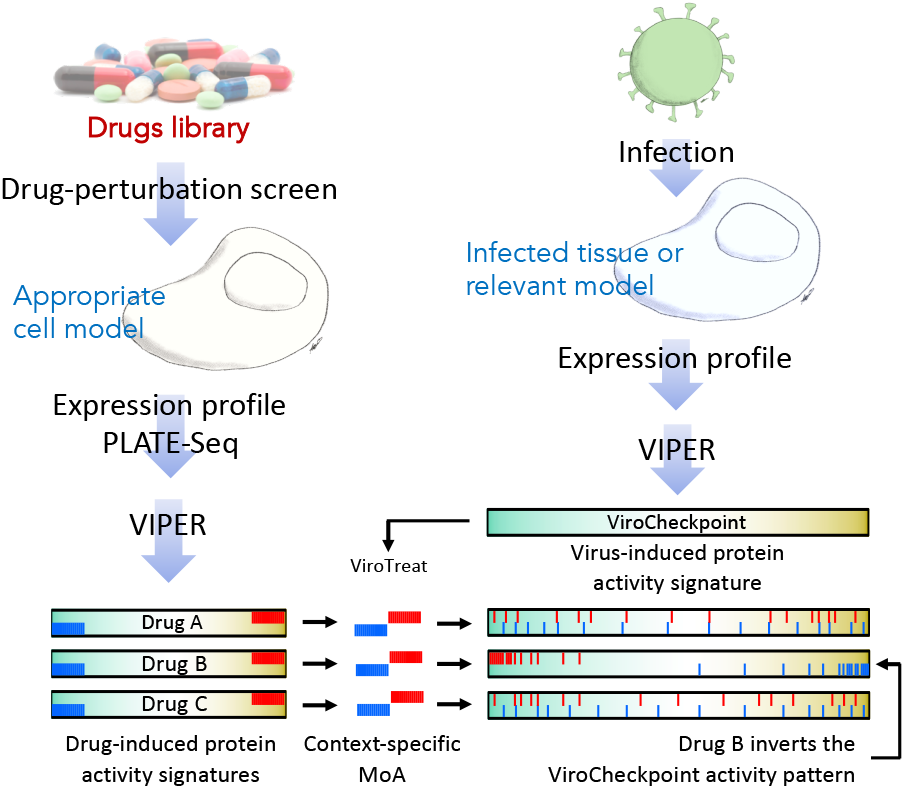
ViroTreat diagram. ViroTreat requires two components: (A) a context-specific drug Mechanism of Action (MoA) database, which is generated by perturbing an appropriate cell model with therapeutically relevant drug concentrations, followed by VIPER analysis of the drug-induced gene expression signatures and identification of the top most differentially active proteins, both activated and inactivated in response to the drug; and (B) the specific virus-induced protein activity signature—where the most differentially active proteins constitute the ViroCheckpoint—dissected by VIPER analysis of a gene expression signature, obtained by comparing an infected tissue or relevant model with non-infected mock controls. ViroTreat then predicts the effect of the drugs on the ViroCheckpoint by matching their MoA with the virus-induced protein activity signature, and quantifies the inverse enrichment using the aREA algorithm. The diagram shows 3 drugs, where only drug B, by activating the host proteins that are being inactivated during virus infection, and inactivating the proteins that are being activated by the virus infection, effectively acts by inverting the ViroCheckpoint activity pattern; and, therefore, would be prioritized as a host cell-targeted antiviral therapeutic option.

ViroTreat requires a tissue-matched drug perturbation database. For this analysis, we had previously generated a collection of RNASeq profiles of NCI-H1793 lung adenocarcinoma cells, at 24h following treatment with a repertoire of 133 FDA approved and 195 late-stage (Phase 2 and 3) drugs—primarily used in or developed for the oncology setting—at their highest subtoxic concentration (48h *IC*_20_) or maximum serum concentration (*C_max_*), whichever is lower. RNASeq data was generated using a fully automated, 96-well based microfluidic technology called PLATE-Seq(55) (Supplementary Table 2). Selection of the NCI-H1793 cell line as an adequate model for the analysis was based on the significant overlap of SARS-CoV infection MR proteins with proteins differentially activated in this cell line (*p* < 10^−28^, 10^−38^, and 10^−24^ at 12h, 24h and 48h after infection, by 1-tail aREA test; see Supplementary Fig. 2). In addition, the main rationale for these assays is the elucidation of protein-level MoA of a drug repertoire and MoA is generally well-recapitulated in lineage matched cells(56).

Using this predictive model, ViroTreat prioritized 44 FDA-approved drugs and 49 investigational compounds in oncology, based on their ability to significantly invert the ViroCheckpoint protein activity signature, at one or more of the 3 evaluated time-points following infection (*p* < 10^−10^, BC; see Supplementary Table 3). Based on this analysis, two FDA-approved drugs—the CDK inhibitor palbociclib and the MEK inhibitor trametinib—and 4 investigational compounds, including three MAP kinase and one AKT/CHEK1 inhibitors, were able to significantly invert the ViroCheckpoint activity at all three time-points (*p* < 10^−10^, BC, Fig. 4a). In addition, six FDA-approved drugs and seven investigational compounds demonstrated the capacity to invert the ViroCheckpoint protein activity pattern at the two earliest time points (12h and 24h, *p* < 10^−10^, BC, Fig. 4a); while two FDA-approved drugs—the ALK and EGFR inhibitors brigatinib and osimertinib—and five investigational compounds were predicted to significantly invert the MR signature identified at later time points (24h and 48h, *p* < 10^−10^, BC, Fig. 4a).

**Fig. 4.**
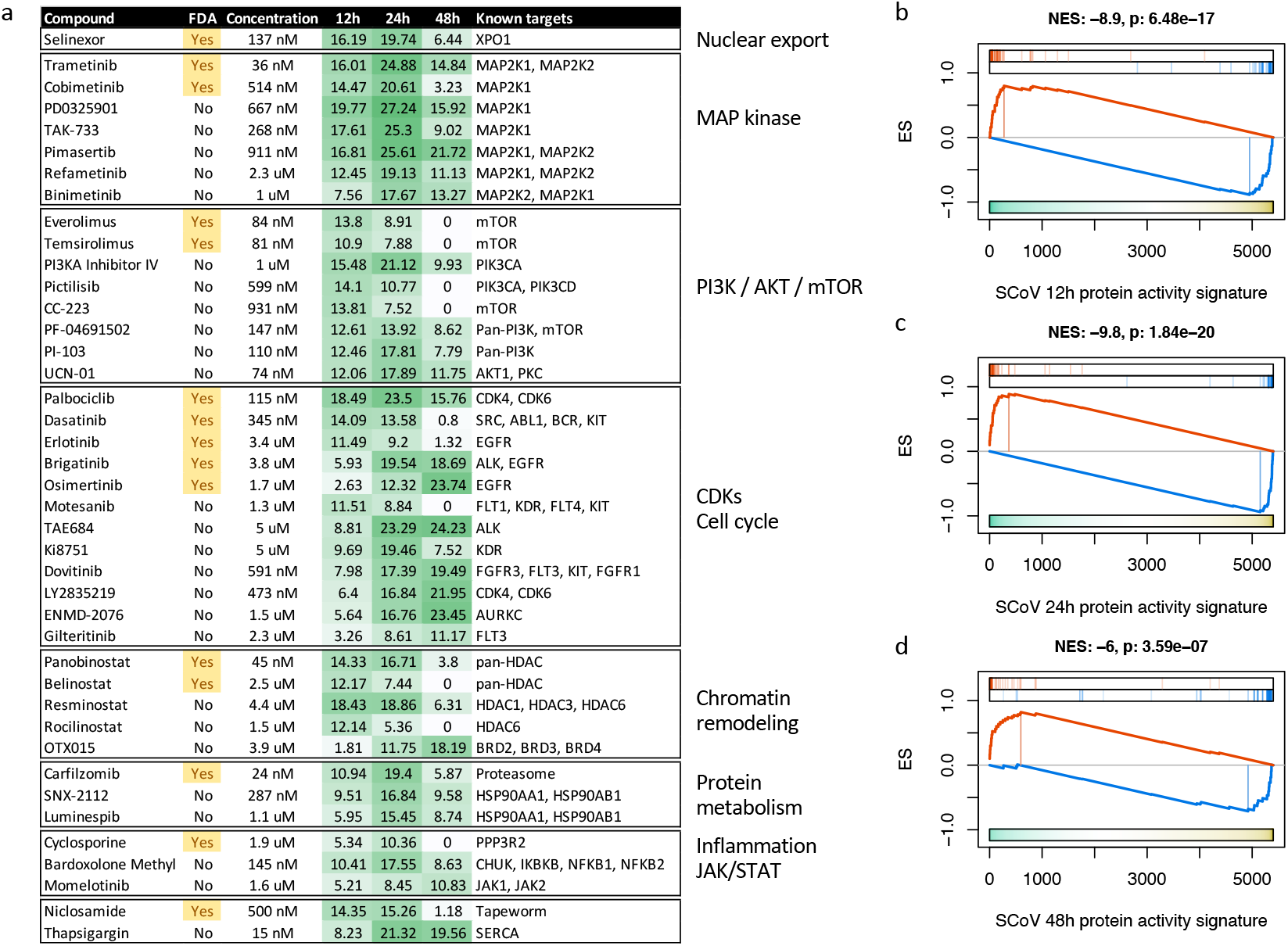
Top drugs and compounds identified by ViroTreat. (a) Table of FDA-approved drugs and investigational compounds identified by ViroTreat as significantly inverting the pattern of activity of the SARS-CoV induced checkpoint (*p* < 10^−10^, BC) for at least one of the three analyzed time points, and being simultaneously significant (*p* < 10^−5^, BC) for at least another time point. The drugs and compounds were organized in blocks according to the biological role or pathway membership of their primary target protein. For each block, the drugs and compounds significant for each time point (*p* < 10^−10^, BC), were sorted by their ViroTreat significant level for 12h, followed by 24h and 48h. FDA-approved drugs were reported prior to investigational compounds. The table also shows the concentration used to perturb NCI-H1793 cells, the ViroTreat significance level, as −log_10_(*p-value*), BC, indicated by the green heatmap, and the primary target for each of the significant drugs and compounds. (b–d) GSEA plots showing the enrichment of the top 25 proteins most activated (red vertical lines), and the top 25 proteins most inactivated (blue vertical lines), in NCI-H1793 cells in response to selinexor perturbation, on the protein activity signatures induced by SARS-CoV infection of Calu-3 cells (x-axis) for 12h (b), 24h (c) and 48h (d). NES and *p-value*, estimated by 2-tail aREA test, are indicated on top of each plot.

Consistent with the pathways enrichment analysis (Fig. 2), several drug families were enriched among the top ViroTreat predictions, including MAP kinases, PI3K/AKT/mTOR, CDK and other cell cycle-related drugs; HDAC and bromodomain protein inhibitors; proteasome and HSP90 inhibitors; and NF-*κ*B and JAK inhibitors (Fig. 4a).

Of special clinical relevance in the context of the COVID-19 pandemic, ViroTreat independently identified the Selective Inhibitor of Nuclear Export (SINE) drug selinexor—FDA-approved for the treatment of relapsed or refractory multiple myeloma—as an extremely potent inverter of SARS-CoV induced ViroCheckpoint activity, in particular, at 12h and 24h time points after infection (*p*_12*h*_ < 10^−16^ and *p*_24*h*_ < 10^−19^, BC, Fig. 4).

## Discussion

ViroTreat presents an application of the extensively validated OncoTreat algorithm for targeting MR proteins driving virus-mediated, reprogrammed cell states induced by viral hijacking of the host cell regulatory machinery. It also provides proof-of-concept of the ability to rapidly prioritize drugs capable of abrogating the reprogrammed, transcriptionally-locked state induced by viral infection, responsible for creating an environment permissive to viral replication and release. Our analysis identified 44 FDA-approved and 49 investigational agents capable of virtually abrogating the MR signature—the ViroCheckpoint protein activity pattern— induced by SARS-CoV infection.

Consistent with the observation that coronaviruses interfere with cell cycle progression to benefit from the physiology of host cells arrested in S phase(26), we show SARS-CoV infection-induced activation of MRs involved in cell cycle progression and DNA repair pathways. Notably, it has been reported previously that coronaviruses inhibit the pRb tumor suppressor protein, inducing infected cell to progress rapidly from *G*_1_ and to arrest the host cell in *S* phase(57). SARS-CoV further favors host cell arrest in *S* phase by inhibiting CDK4 and CDK6 kinase activity(58). We also observed activation of PI3K/AKT/mTOR pathway proteins, suggesting that SARS-CoV—similar to other viruses(59), including +ssRNA viruses like chikungunya(60), hepatitis C(61), west nile(62) and dengue(63), as well as other RNA respiratory viruses like influenza(64) and the respiratory syncytial virus(65)—might subvert mTOR pathway activity. Indeed, temporal kinome analysis of human hepatocytes infected with MERS-CoV had previously revealed changes in MAPK and PI3K/AKT/mTOR pathways(20). Finally, we observed activation of proteins involved in innate immunity, including interferon response and pro-inflammatory pathways, which have been also previously described for coronaviruses(26).

While formal experimental validation is still required, there are several positive indications this approach may be effective. Specifically, drugs for SARS-CoV most highly prioritized by ViroTreat were highly consistent, at least based on their primary target proteins, with biological programs and pathways known to be modulated by coronavirus infection(26, 66). Notably, in this regard, cell cycle progression/proliferation, PI3K/AKT/mTOR, innate immunity and inflammation are well represented among the primary target proteins for those pharmacologic agents strongly predicted by ViroTreat to possess host cell-targeted, antiviral effects.

A literature search revealed that many of the oncology drugs and compounds identified by ViroTreat have been considered previously for their potential antiviral effects. For instance, the MAPK inhibitor trametinib, one of the top ViroTreat hits for SARS-CoV, was shown to inhibit MERS-CoV replication *in vitro*(5, 20), as well as influenza A virus both *in vitro* and *in vivo*(67). Similarly, everolimus, an mTOR inhibitor identified by ViroTreat, has also been shown to inhibit MERS-CoV(5, 20) and cytomegalovirus(68) replication *in vitro*, as well as to reduce incidence of cytomegalovirus infections following kidney transplant(69). Among tyrosine kinase inhibitors identified by ViroTreat, dasatinib was previously described to inhibit MERS-CoV(5, 19) and HIV-1(70) replication *in vitro*; while erlotinib was shown to inhibit dengue(71), hepatitis C(72) and ebola(73) replication. The HSP90 inhibitors SNX-2112 and luminespib, as well as the sarco/endoplasmatic reticulum Ca^2+^ ATPase inhibitor thapsigargin, all identified by ViroTreat as inverters of the SARS-CoV induced checkpoint, have been shown to inhibit herpes simplex(74), chikungunya(75), foot and mouth disease virus(76), respiratory syncytial virus(77), rhinovirus(78) and hepatitis A virus replication(79).

Finally, ViroTreat independently identified the SINE drug molecule selinexor—an FDA-approved agent for the treatment of relapsed or refractory multiple myeloma—as an extremely potent inverter of SARS-CoV-induced ViroCheckpoint activity. Selinexor is a potent and highly-specific inhibitor of XPO1 activity, which leads to nuclear retention of its cargo proteins containing leucine rich Nuclear Export Signals. Based on experimental studies performed by Karyopharm Therapeutics Inc., low Selinexor concentrations (*leq* 100 nM) inhibited viral replication by 90% in green monkey kidney Vero cells infected with SARS-CoV-2(80). As a result of these observations and data, which are consistent with the ViroTreat prioritization of selinexor we report in this study, a randomized, placebo-controlled Phase 2 clinical study (NCT04355676 and NCT04349098), evaluating low dose oral selinexor in hospitalized patients with severe COVID-19 has been initiated and is currently enrolling patients, with results anticipated to be reported by August 31^st^, 2020(80).

This analysis has several limitations that partially restrict its value as proof of concept. Specifically, infection was conducted in a cancer cell line, rather than in a more physiologically relevant context, such as in primary bronchial or alveolar epithelial cells. In addition, drug perturbations were also performed in a cancer cell line context, thus potentially introducing undesired confounding factors, even though use of mock controls for the infection, and vehicle control for the drug perturbations, from the same cancer cell line should have eliminated most of the cancer-related bias and cell line idiosyncrasies. As a result, extrapolation of this approach to the clinic may be limited by the following assumptions: (a) that the host cell regulatory checkpoint hijacked by the virus is conserved between the Calu-3 adenocarcinoma cell line and the normal alveolar or bronchial epithelial cells *in vivo*; and (2) that the drugs’ and compounds’ MoA is conserved between the NCI-H1793 lung adenocarcinoma cells and normal lung epithelial cells *in vivo*. Moreover, while for the generation of the perturbational data and the context-specific MoA database we used subtoxic drug concentrations that, in most cases, were well below the maximum tolerated dose for all drugs and compounds, the relevant pharmacologic concentration for their deployment as antiviral therapy may be much lower than the original recommended concentration for their use as anti-cancer drugs.

Further research is necessary to benchmark the ViroTreat approach. Specifically, better reporters of SARS-CoV infection should be established, ideally directly from nasopharyngeal swabs or bronchial lavage of SARS-CoV patients. More relevant to the current pandemic, such samples are starting to emerge from COVID-19 patients and may lead to elucidation of critical entry points for COVID-19 therapeutic intervention. Similarly, drug profiles should be generated in a more physiologic context, including primary airway epithelial cells. It is also important to establish whether virus-induced transcriptional lock states are similar across all cell and tissue contexts infected by the virus, or whether the hijacked states are cell context-specific. Finally, appropriate environments for *in vitro* and *in vivo* validation of prioritized drugs should be developed(56).

To our knowledge, this is the first time a virus-induced MR module (i.e., the ViroCheckpoint) is proposed as a pharmacological target to abrogate the virus’s ability to hijack the cellular machinery of host cells, a strategy that coronaviruses are known to employ to prime the host cell environment so it is amenable to viral replication and release(26). In addition, ViroTreat represents a unique method for the systematic and quantitative prioritization of mechanism-based, host-directed drugs capable of abrogating this critical, and previously unaddressed component of viral infection. If effectively validated, this approach presents several advantages: First, ViroTreat is tailored to target the entire repertoire of host proteins hijacked by the virus to create a permissive environment, rather than a single host or viral protein. As such, we anticipate drugs identified by ViroTreat to have more universal applications, including being effective against a broader viral repertoire, while also being more effective at eluding virus adaptation mechanisms arising from rapid mutation under drug selection stress. Indeed, drug-mediated reprogramming of host cell to a transcriptional state that confers resistance against coronavirus-induced reprogramming presents the opportunity to identify drugs that are potentially effective for a broader class of viruses, as long as they share similar pathobiological strategies for host cell takeover. Second, the ViroTreat analysis can be performed expeditiously—as soon as the ViroCheckpoint signature of a novel virus becomes available. Therefore, this methodology is especially well-suited to the urgency characteristic of epidemics and pandemics.

Developing effective treatments for respiratory tract infections—i.e., those that reduce such hard end points as hospitalization, need for mechanical ventilation, and mortality—exclusively through direct viral targeting has been historically challenging. Drugs identified specifically for host cell-targeting have the potential therapeutic advantage of acting in a mechanistically complementary—even synergistic—way with readily available antivirals, thereby suggesting roadmaps for developing and testing combination regimens that may mitigate viral replication by acting upon the infected system as a whole. Such multi-mechanistic pharmacologic approaches targeting both the virus and host cell proteins that are critically dysregulated as a result of viral infection may be required to optimize clinical outcomes, especially in challenging and vulnerable patients exposed to lethal pathogens with high virulence and viral load.

## ACKNOWLEDGEMENTS

We thank Christopher Walker for reviewing selinexor data accuracy and Tatiana Alvarez for original artwork. This research was supported by the following NIH grants to Andrea Califano: R35 CA197745 (Outstanding Investigator Award); U01 CA217858 (Cancer Target Discovery and Development); S10 OD012351 and S10 OD021764 (Shared Instrument Grants).

## Author Contributions

Conceptualization and Methodology, P.L., A.C. and M.J.A.; Investigation, P.L., X.S., G.B. and M.J.A.; Formal Analysis, P.L., X.S., Y.S., E.F.D. and M.J.A.; Experimental execution and data generation: C.K., R.R. and S.P.; Original Draft, G.B., A.C. and G.B.; Writing – Review and Editing, P.L., G.B., A.C. and M.J.A.

## Methods

### Cell lines

NCI-H1793 cells were obtained from ATCC (CRL-5896), mycoplasm tested and maintained in DMEM:F12 medium supplemented with 5 *μ*g/ml insulin, 10 *μ*g/ml transferrin, 30 nM sodium selenite, 10 nM *β*-estradiol, 4.5 mM L-glutamine and 5% fetal bovine serum. Cells were grown in a humidified incubator at 37°C and 5% CO_2_.

### Lung epithelium context-specific drug mechanism of action database

The drug-perturbation dataset was generated as follows. First, the *ED*_20_ for each of the 133 FDA-approved drugs and 195 investigational compounds in oncology was estimated in NCI-H1793 cells by performing 10-point dose-response curves in triplicate, using total ATP content as read-out. Briefly, 2,000 cells per well were plated in 384-well plates. Small-molecule compounds were added with a 96-well pin-tool head 12h after cell plating. Viable cells were quantified 48h later by ATP assay (CellTiterGlo, Promega). Relative cell viability was computed using matched DMSO control wells as reference. *ED*_20_ was estimated by fitting a four-parameter sigmoid model to the titration results. NCI-H1793 cells, plated in 384-well plates, were then perturbed with a library of 328 FDA-approved drugs and small-molecule compounds at their corresponding *ED*_20_ concentration. Cells were lysed at 24h after small-molecule compound perturbation and the transcriptome was profiled by PLATE-Seq(55). RNA-Seq reads were mapped to the human reference genome assembly 38 using the STAR aligner(81). Expression data were then normalized by equivariance transformation, based on the negative binomial distribution with the DESeq R-system package (Bioconductor(82)). At least two replicates for each condition were obtained. Differential gene expression signatures were computed by comparing each condition with plate-matched vehicle control samples using a moderated Student’s t-test as implemented in the limma package from Bioconductor(83). Individual gene expression signatures were then transformed into protein activity signatures with the VIPER algorithm(37), based on the a lung adenocarcinoma context-specific regulatory network available from the aracne.networks package from Bioconductor.

### Computational analysis

Enrichment of gene-sets for biological hallmarks was performed using Gene Set Enrichment Analysis(52) with the Molecular Signatures Database MSigDB v7.1(53). Enrichment analysis for virus-interacting host proteins (PPI) on SARS-CoV induced protein activity signatures, as well as the OncoMatch(56) analysis to assess the conservation of the virus-induced MR protein activity on NCI-H1793 lung adenocarcinoma cells were performed with the aREA algorithm(37).

### ViroTreat analysis

ViroTreat was performed by computing the enrichment of the top/bottom 50 most differentially active proteins in response to drug perturbation—the context-specific mechanism of action—on the virus-induced protein activity signature using the aREA algorithm(37). P-values for significantly negative enrichment were estimated using 1-tail aREA analysis, and multiple hypothesis testing was controlled by the Bonferroni’s correction.

### Code availability

All the code used in this work is freely available for research purposes. VIPER and aREA algorithms are part of the “viper” R-system’s package available from Bioconductor. The lung adenocarcinoma context-specific interactome is available as part of the “aracne.networks” R-system’s package from Bioconductor.

## Supplementary Figures and Tables

**Supplementary Figure S1.**
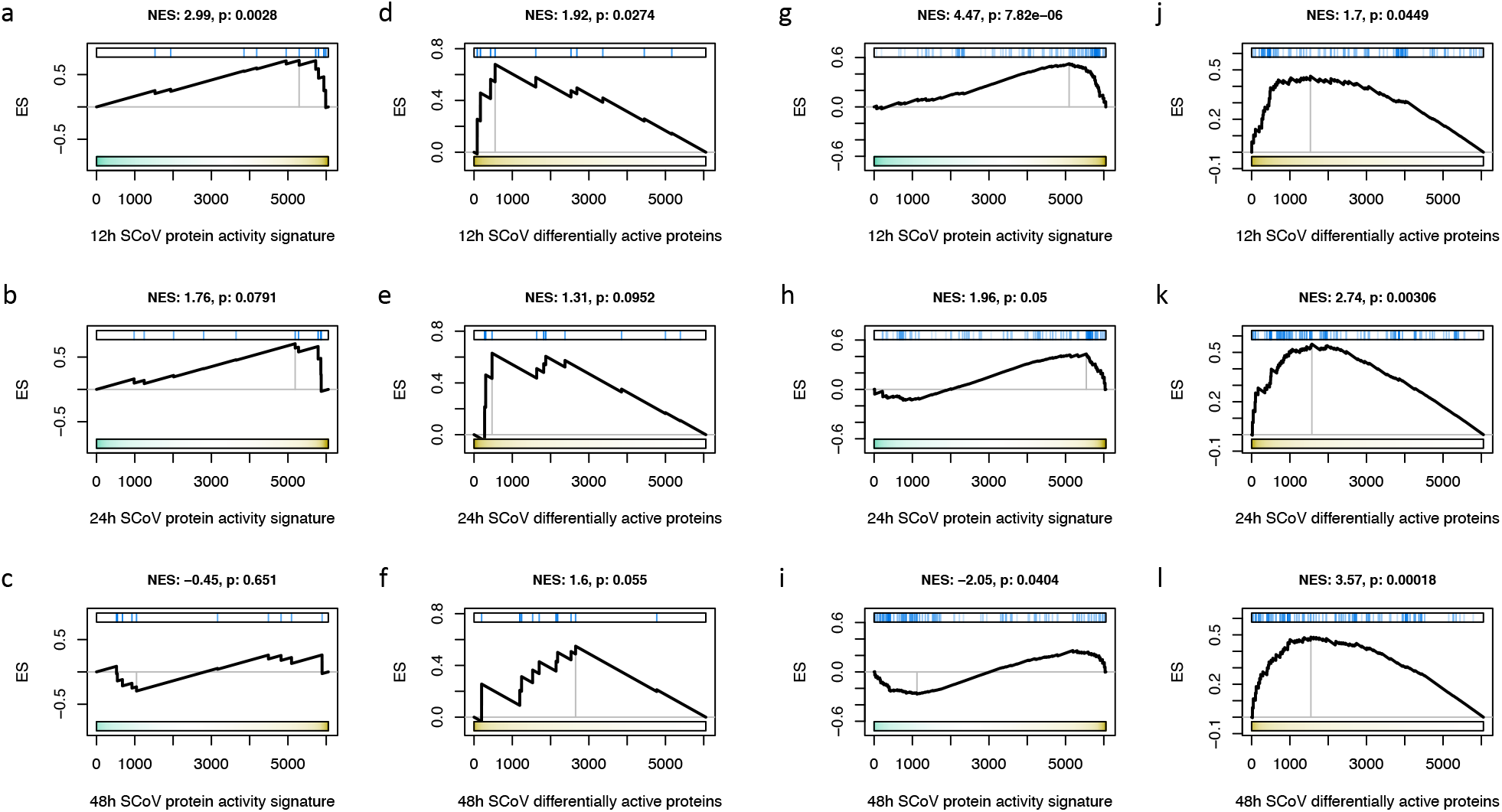
Enrichment of SARS-CoV- and SARS-CoV2-interacting host proteins among the most differentially active proteins after SARS-CoV infection. (a–f) Enrichment of 12 SARS-CoV-interacting host proteins, or (g–l) 89 SARS-CoV2-interacting proteins on SARS-CoV induced protein activity signatures at 12h (a, d, g and j), 24h (b, e, h and k) and 48h (c, f, i and l) after infection. GSEA plots show the enrichment score (y-axis) and the SARS-CoV induced protein activity signature (x-axis), where 6,054 regulatory proteins were rank-sorted from the one showing the strongest inactivation (left) to the one showing the strongest activation (right) in response to SARS-CoV infection (a–c and g–i); or where the regulatory proteins were sorted from the most differentially active (left) to the least differentially active (right) after SARS-CoV infection. NES and *p-value* were estimated by 2-tail aREA test and shown on top of each plot.

**Supplementary Figure S2.**
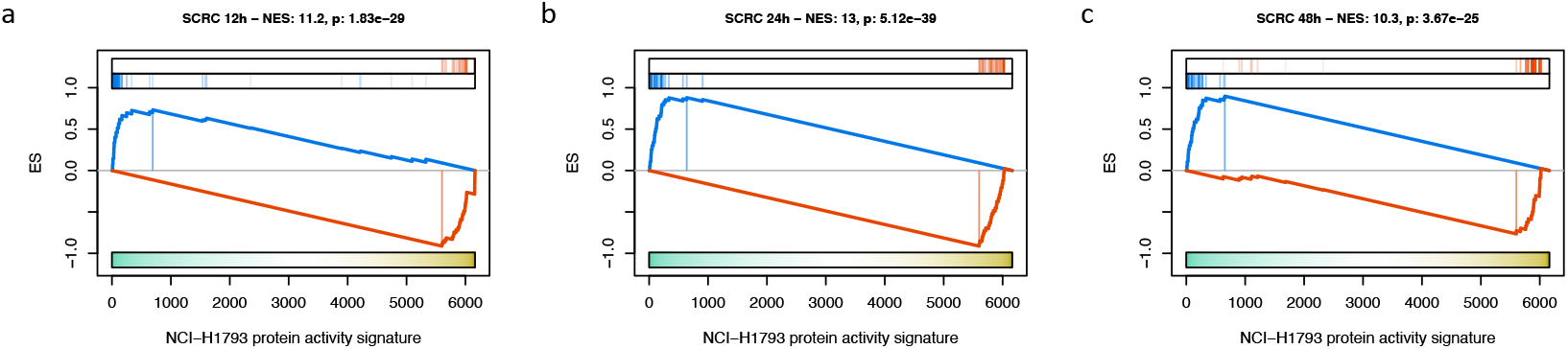
Conservation of the SARS-CoV induced checkpoint in NCI-H1793 cells. GSEA plots for the enrichment of the top 25 most activated proteins (red vertical lines), and top 25 most inactivated proteins (blue vertical lines) by SARS-CoV infection at 12h (a), 24h (b) and 48h (c) after infection. The x-axis shows 6,054 proteins rank-sorted from the most inactivated ones (left) to the most activated ones (right) in NCI-H1793 cells when compared against 86 non-small cell lung cancer cell lines. The y-axis shows the GSEA enrichment score. NES and *p-value*, estimated by 2-tail aREA test, are indicated on top of each plot.

**Supplementary Table 1.**
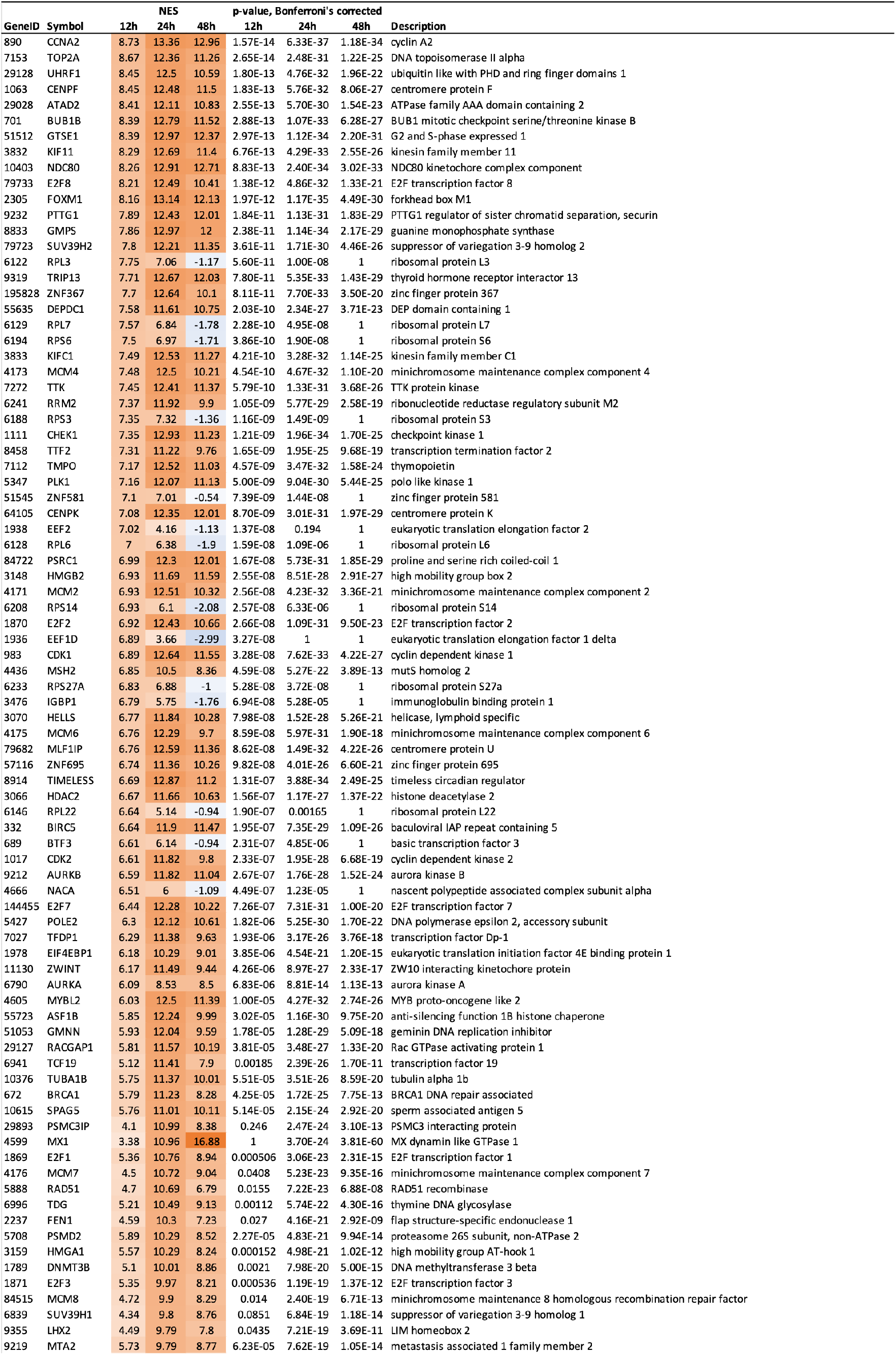
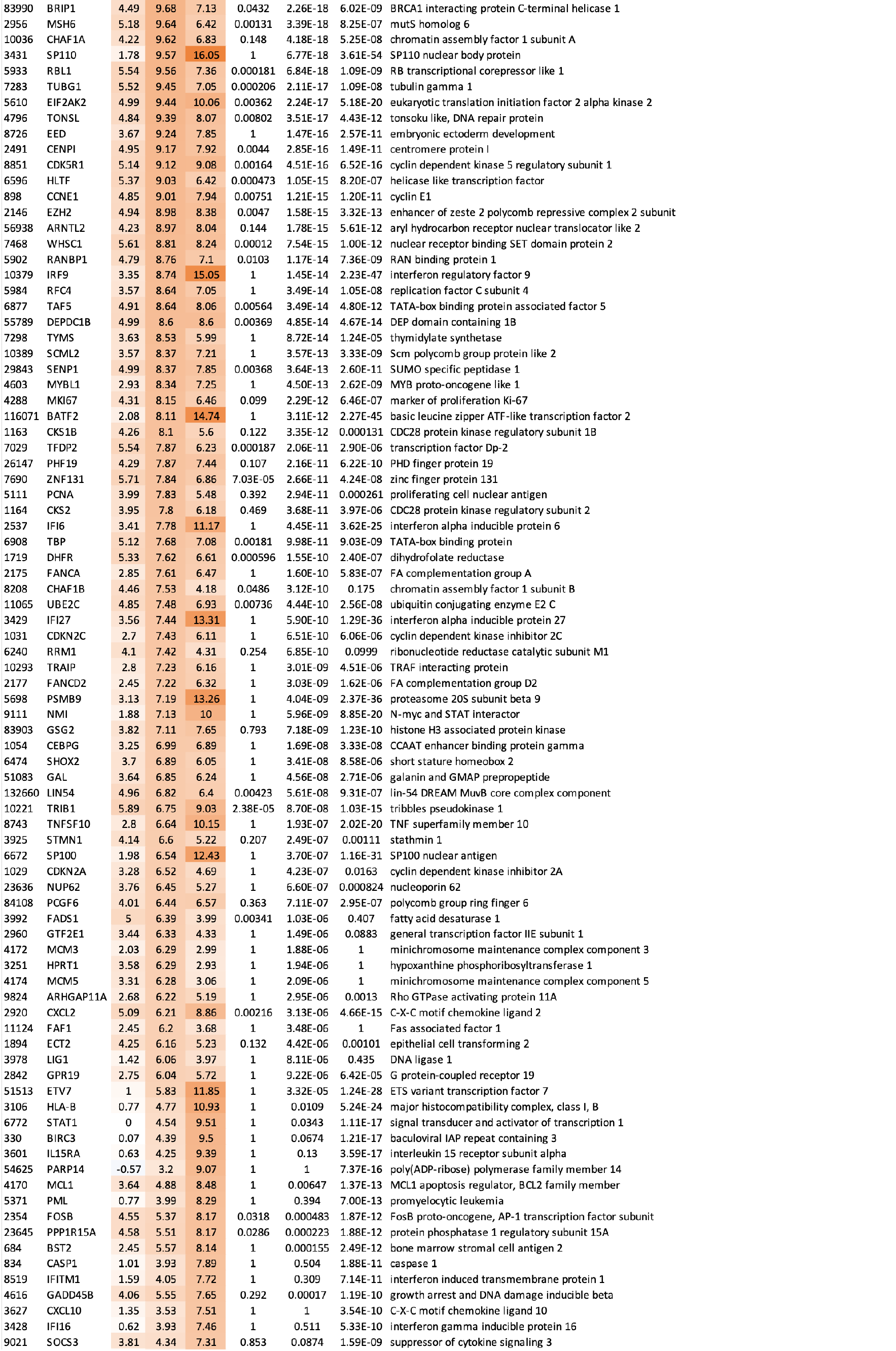
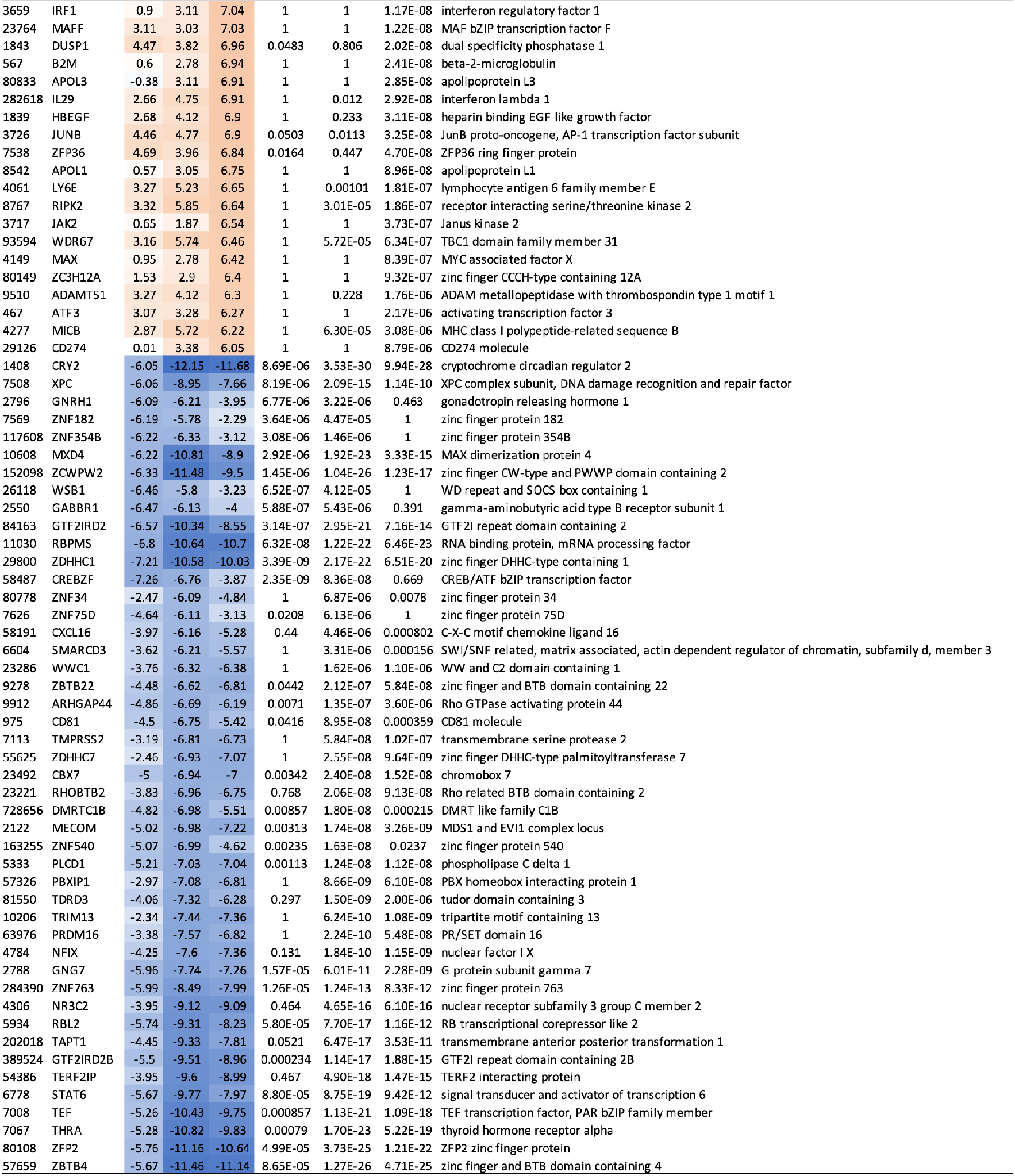
Proteins differentially active in response to SARS-CoV infection. Shown are 236 proteins differentially active (*p* < 10^−5^, BC, 2-tail aREA test) at any of the three evaluated time points. The table includes the EntrezID, and symbol of the genes coding for the differentially active proteins, the VIPER-inferred NES and Bonferroni’s corrected *p-value*.

**Supplementary Table 2.**
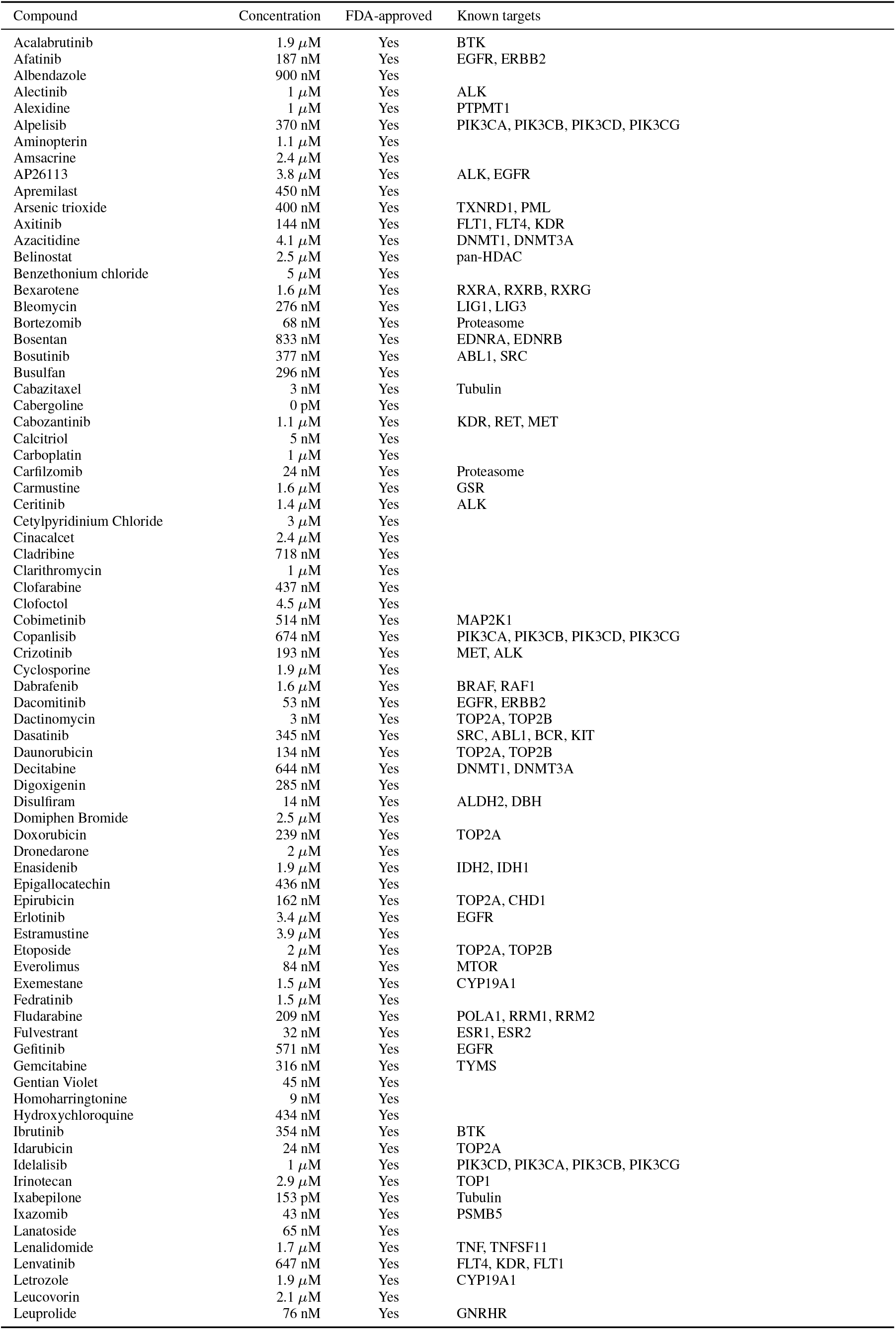
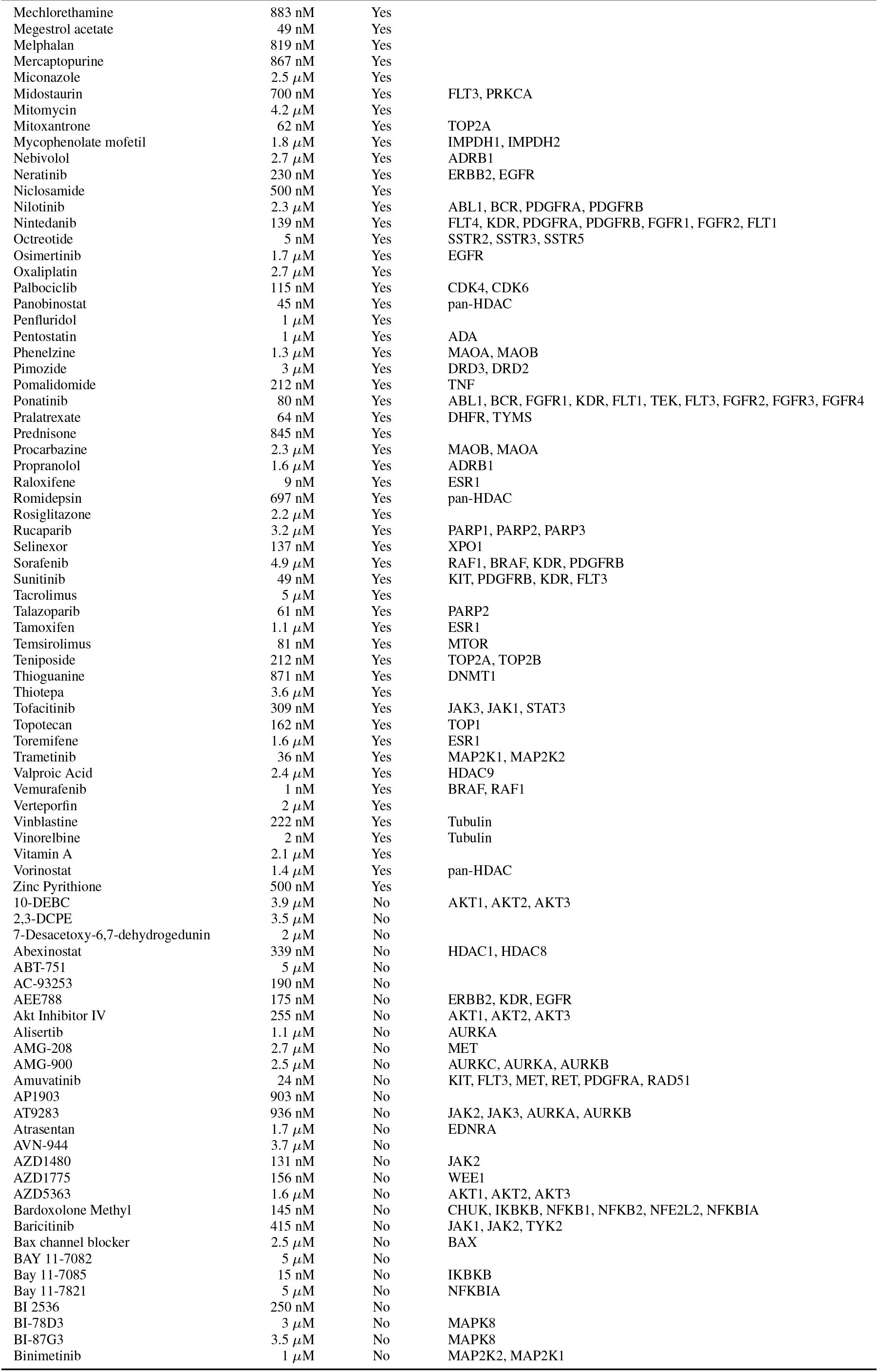
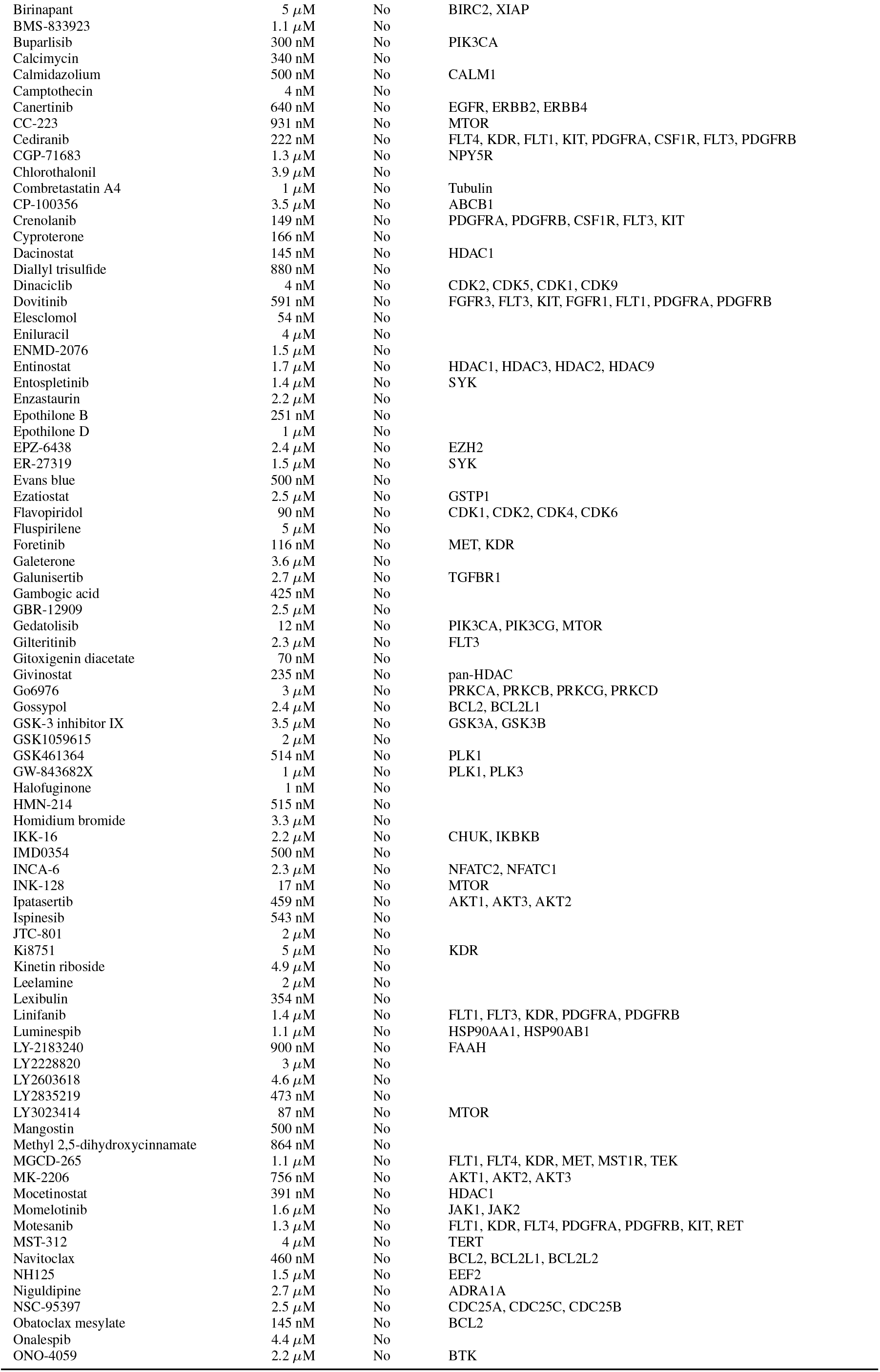
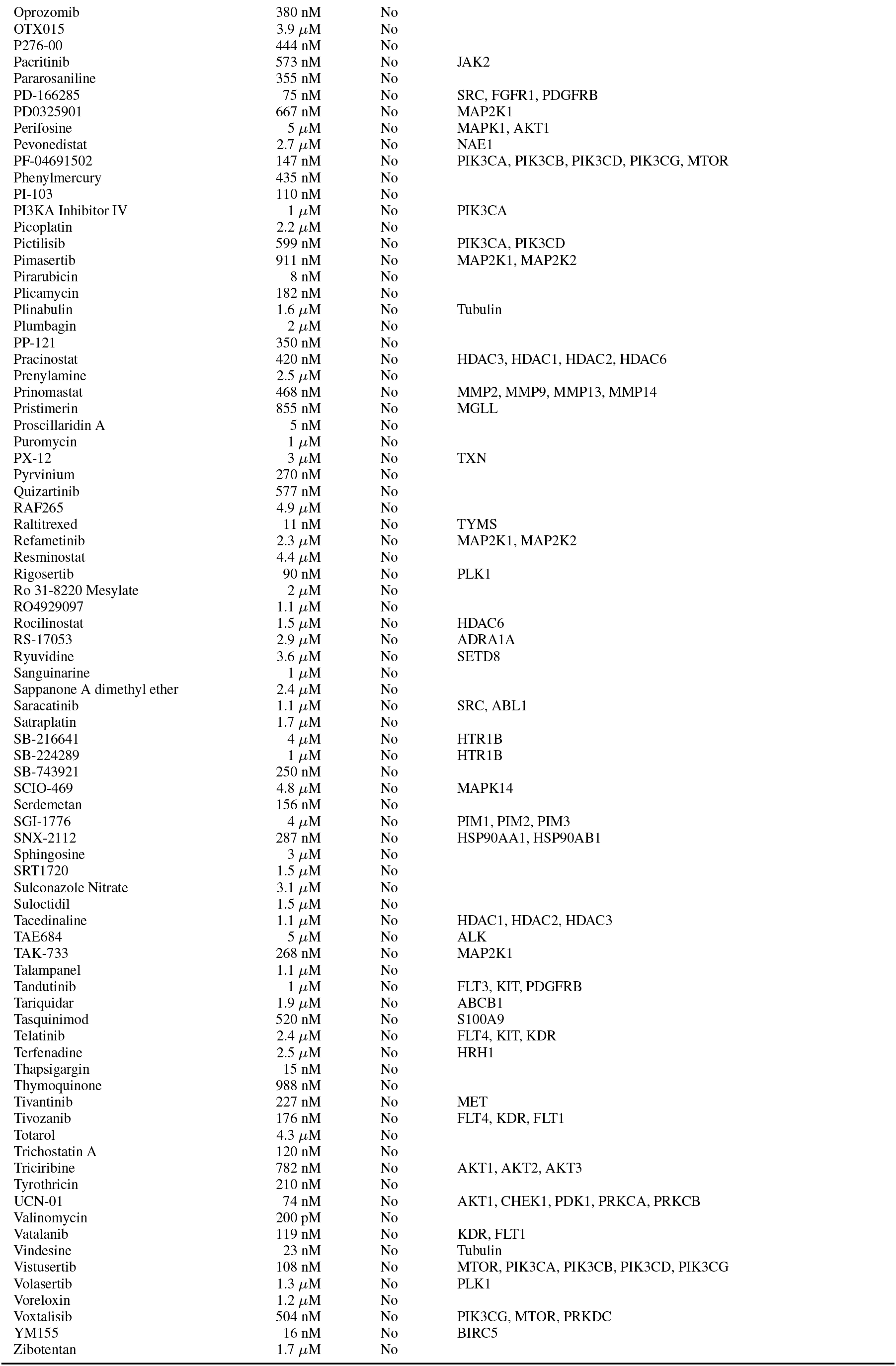
FDA-approved drugs and late-stage (phase 2 and 3) investigational compounds in oncology covered by the lung epithelium context-specific MoA database. The table lists the drug/compound name, concentration used to perturb NCI-H1793 lung adenocarcinoma cells, FDA-approval status known primary targets.

**Supplementary Table 3.**
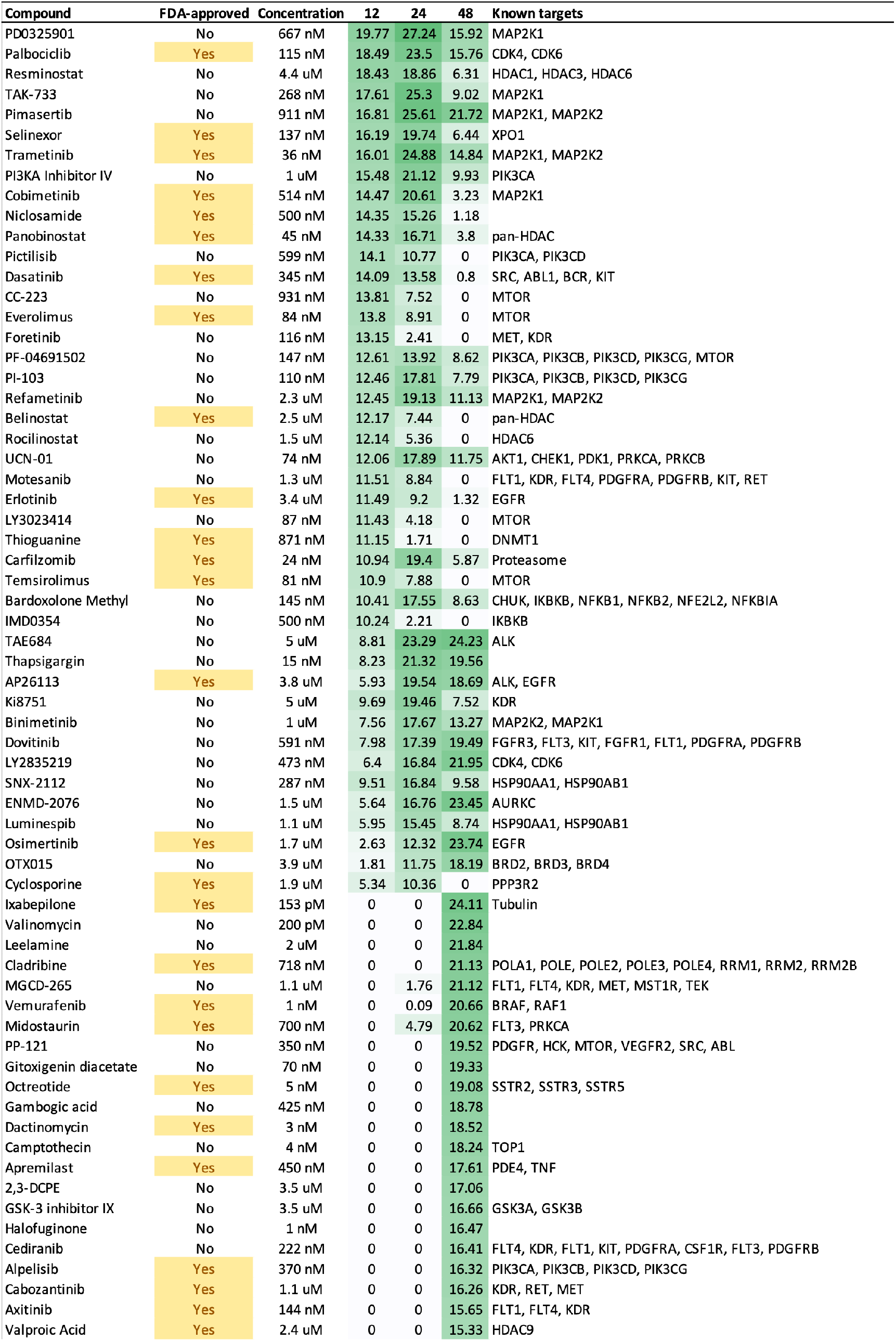
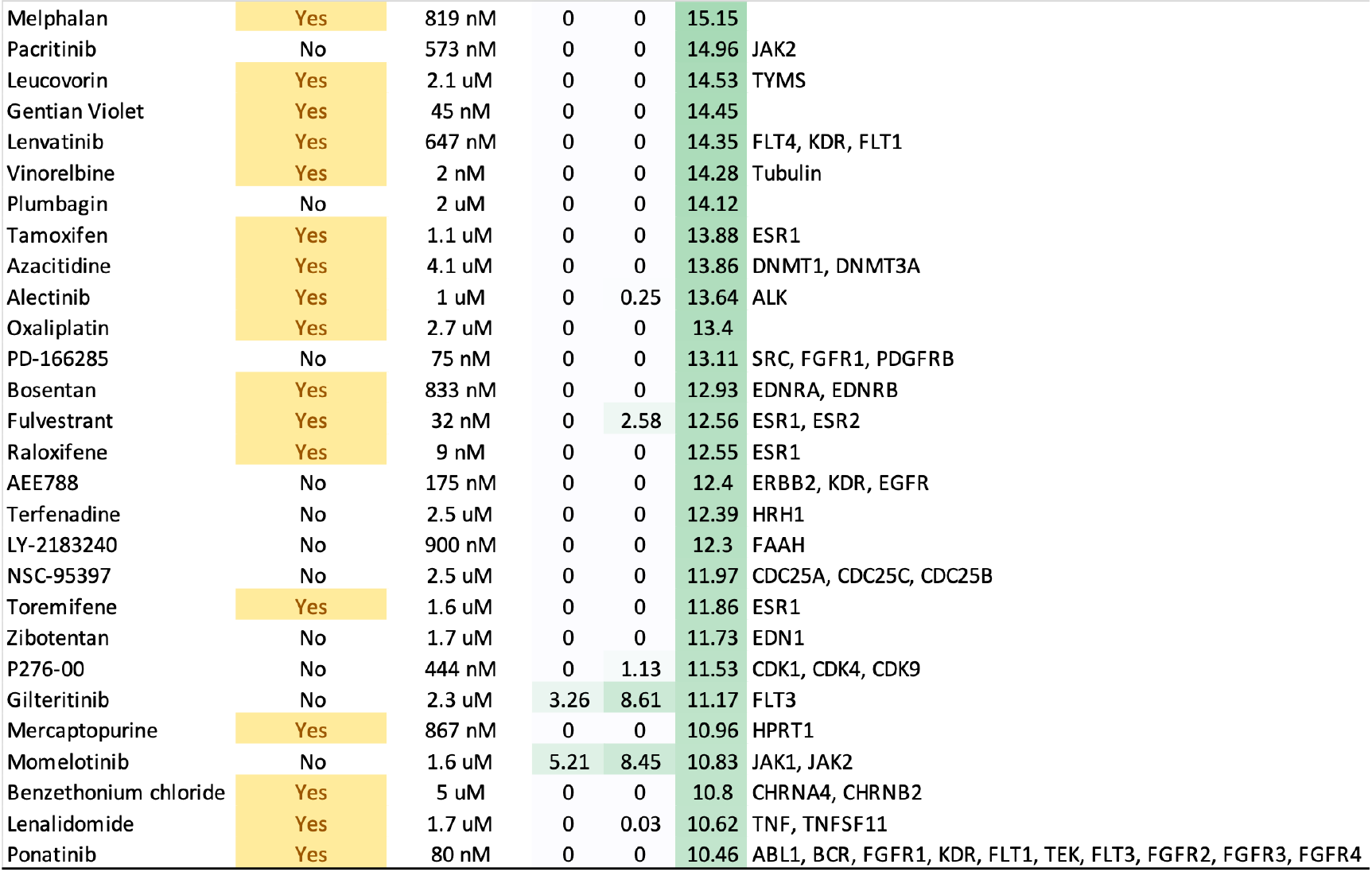
FDA-approved drugs and investigational compounds identified by ViroTreat as significantly inverting the SARS-CoV ViroCheckpoint (*p* < 10^−10^, BC, 1-tail aREA test). The drugs/compounds were sorted according to ViroTreat-inferred statistical significance as inverters of SARS-CoV 12h-, 24h- and 48h-ViroCheckpoints. The table lists the drug/compound name, FDA-approval status, concentration used to perturb the NCI-H1793 lung adenocarcinoma cells, ViroTreat-estimated statistical significance—expressed as −log_10_(*p-value*)—and know primary targets.

